# Heterozygous mutations in *p97* and resistance to p97 inhibitors

**DOI:** 10.1101/380964

**Authors:** Prabhakar Bastola, Kay Minn, Jeremy Chien

**Affiliations:** Division of Molecular Medicine, Department of Internal Medicine, University of New Mexico, Health Sciences Center, Albuquerque, New Mexico 87131, USA

## Abstract

In recent years, multiple studies including ours have reported on the mechanism of resistance towards p97 inhibitors. While all these studies outline target alteration via mutations in p97 as the primary mechanism of resistance, discrepancies persist in the current literature due to the occurrence of both heterozygous and homozygous mutations when using HCT116 cells. Here, we report a pre-existing heterozygous frameshift mutation at codon 616 (N616fs*) in one of the *p97* alleles in HCT116 cells and show that this mutant allele is subjected to nonsense-mediated decay. Furthermore, we independently generated p97 inhibitor (CB-5083) resistant HCT116 cells, and we observed a single heterozygous mutation at codon 526 (L526S) in genomic DNA sequencing but a homozygous L526S mutation in complementary DNA sequencing, indicating that the missense mutation (L526S) occurs in the allele that does not harbor the frameshift N616fs* mutation. Our results underscore the importance of performing simultaneous genomic and complementary DNA sequencing when confirming mutations in *p97*.

## Introduction

With p97 inhibitor CB-5083 entering the realm of clinical trials, multiple studies have evaluated the resistance mechanisms towards different classes of p97 inhibitors. Her *et al.* reported that a single heterozygous missense mutation in *p97* (A530T) is sufficient to produce resistance to the p97 inhibitor NMS-873 in HCT116 cells [1]. Likewise, Anderson *et al.* reported that single homozygous missense mutations in *p97* are sufficient to produce resistance to the p97 inhibitor CB-5083 in HCT116 cells [2]. Lastly, we recently reported unique patterns of coselected mutations with heterozygous missense mutations in one *p97* allele and heterozygous nonsense mutations in the other *p97* allele resulting in resistance to both p97 inhibitors CB-5083 and NMS-873 in OVSAHO cells [3, 4]. While results from Anderson *et al.* show that homozygous mutations in p97 is necessary for resistance towards CB-5083, the conclusion from Her *et al.* is that a single heterozygous mutation in *p97* is sufficient for resistance towards NMS-873. Based on these discrepancies, Her *et al*. speculated that the difference in homozygous and heterozygous mutations associated with resistance to p97 inhibitors may reflect the potential difference in mechanisms of action between CB-5083 and NMS-873. While the above conclusions could help explain the differences in missense mutations observed in these two studies, their conclusion about the sufficiency of a single heterozygous mutation in *p97* for resistance to NMS-873 should be reconsidered in light of the fact that HCT116 cells harbor a pre-existing frameshift mutation in one allele of *p97.* Here, we report a pre-existing heterozygous frameshift mutation in HCT116 cells. Our results help to explain the discrepancies between these studies as well as highlight the importance of sequencing both the genomic DNA and the complementary DNA when evaluating mutations in *p97* associated with resistance to p97 inhibitors.

## Results and Discussion

Our analysis of HCT116 exome sequencing data from the Cancer Cell Line Encyclopedia (CCLE) indicates HCT116 cells harbor a heterozygous frameshift mutation at codon 616 (N616fs*) (Figure 1A). In total, we found 98 cancer cell lines that harbored mutation in p97 (gene, VCP) among which N616fs* was the most frequently reported mutation in these samples (Supplementary 1). N616fs* is also the most frequently reported mutation in TCGA tumor sample (Supplementary 2). To confirm N616fs* in HCT116 cells, we performed Sanger sequencing of genomic DNA (gDNA) flanking exon 14 of *p97* that contains codon 616 sequences. The Sanger sequencing results confirm that HCT116 cells harbor a single-base deletion in codon 616 that causes a frameshift mutation (Figure 1B). Given that the frameshift mutation introduces a premature stop codon (N616fs*), we suspected that transcripts from the mutated allele will undergo nonsense-mediated decay (NMD). To confirm this possibility, we performed Sanger sequencing of complementary DNA (cDNA) of *p97* from HCT116. Not surprisingly, we did not observe the frameshift mutation at codon 616 in the cDNA sequencing analysis (Figure 1B), confirming that transcripts from the mutated allele are subjected to NMD. This observation is important because prior studies by Her *et al.* and Anderson *et al.* used HCT116 to generate clones that are resistant to VCP inhibitors. Both studies reported mutations in p97 contributing to resistance to VCP inhibitors CB-5083 and NMS-873. While Anderson *et al.* reported that their mutations in *p97* are homozygous, Her et al. reported their mutation in p97 is heterozygous. Her *et al.* concluded that heterozygous mutation in *p97* was sufficient for resistance to NMS-873. These conclusions should now be interpreted with caution given that HCT116 cells harbor a pre-existing truncation mutation that results in NMD.

**Figure 1:**
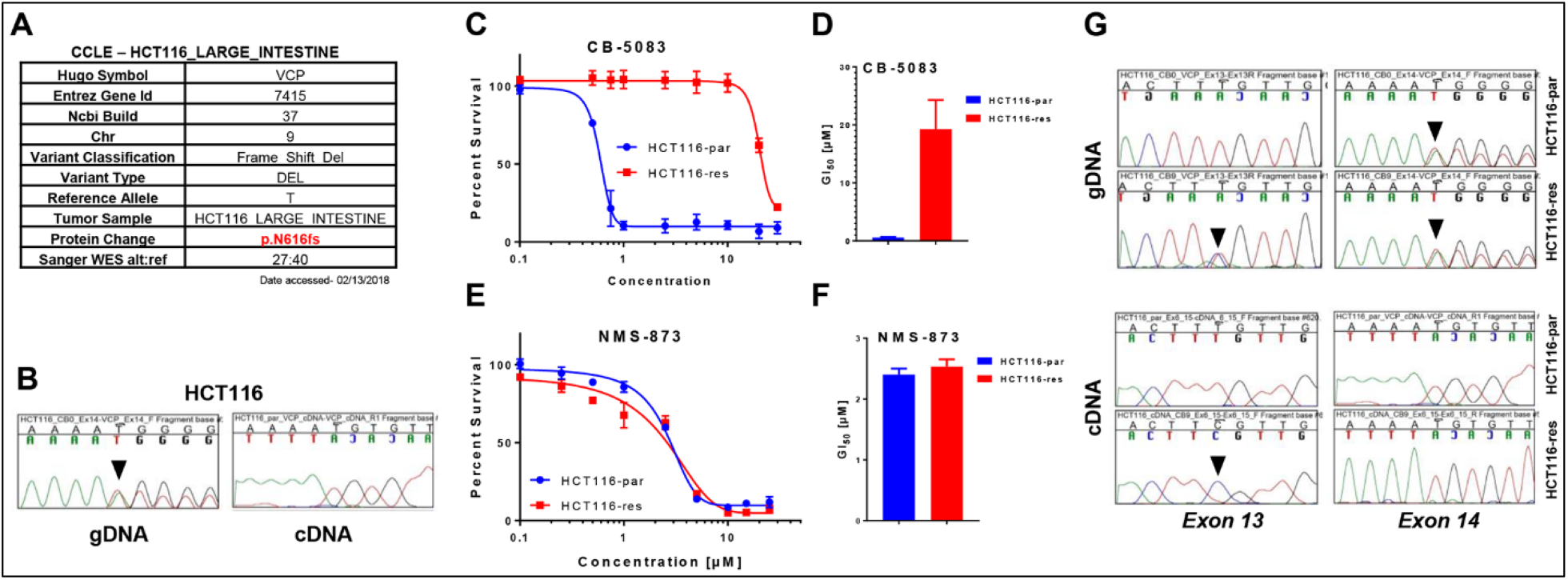
HCT116-resistant cells harbor a heterozygous missense mutation and a heterozygous frameshift mutation in the p97 genomic DNA. A) Analysis of HCT116 exome sequencing results from the Cancer Cell Line Encyclopedia (CCLE). B) Chromatograms of the genomic DNA (gDNA) and complementary DNA (cDNA) sequences displaying partial sequences of p97 exon 16 in parental HCT116 cells. C) Parental HCT116 (HCT116-par) and CB-5083-resistant HCT116 (HCT116-res) cells were incubated with different doses of CB-5083 between 0.1 μM and 30 μM for 72 hours. Dose-response curves were generated with the Graph-Pad Prism using four parameters nonlinear regression and the curves were constrained at the top (100%) and the bottom (>0%). Each point in the dose-response curve represents Mean ± SEM from three technical replicates. D) The bar-graph represents Mean GI50 + SEM obtained from three biological replicates. E&F) Similar to C&D, but treated with increasing doses of NMS-873 between 0.1 μM and 25 μM for 72 hours. G) Chromatograms of the gDNA and cDNA sequences displaying specific regions of *p97* in the HCT116-par and the HCT116-res cells. Black arrows point to the observed mutations in *p97.*

To provide additional evidence that mutated allele is not subjected to additional mutational selection pressure under drug treatment, we independently generated CB-5083 resistant cells in HCT116 (HCT116-res). HCT116-res cells displayed approximately 34-fold resistance towards CB-5083 (Figure 1C & 1D) while displaying no cross-resistance towards NMS-873 (Figure 1E & 1F). Subsequently, we sequenced the genomic DNA and complementary DNA to identify potential mutations in *p97.* In HCT116-res cells, we identified a single heterozygous mutation (A>C) at codon 526 (L526S) in gDNA sequencing but observed a homozygous mutation (A>C) at codon 526 (L526S) in cDNA sequencing (Figure 1G). These results suggest that the L526S mutation occurs in the allele that does not have the pre-existing N616fs* mutation. These results also suggest that the wild-type allele is subjected to a selection pressure under drug treatment in HCT116 cells.

Based on these results, it is more than likely that the single heterozygous missense mutation (A530T) reported by the Her *et al.* resides on the allele that does not have the pre-existing N616fs* mutation and that their resistant cells express solely the mutant p97 proteins. Authors should confirm the exclusive mutations in two *p97* alleles by performing both complementary DNA sequencing and genomic DNA sequencing of their resistant cells. HCT116 cells harbor deficiency in mismatch repair [5] as well as display low expression of P-glycoprotein [6], which makes it an ideal cell line to screen for drug-resistant clones. It is therefore not surprising that HCT116 cells were used to screen for drug resistance. While Anderson *et al.* sequenced the cDNA to identify homozygous mutations in *p97,* Her *et al.* sequenced the gDNA to observe the heterozygous mutation (A530T). By sequencing *p97* at the gDNA and cDNA level, we were able to resolve the conflicting results obtained from these studies. Given that HCT116 cells harbor pre-existing N616fs* mutation that is subjected to NMD, reported studies including ours do not provide conclusive evidence that a single heterozygous mutation in *p97* is sufficient to produce resistance to p97 inhibitors. Given that the functional unit of p97 is a homo-hexamer, a heterozygous mutation in *p97* is expected to produce hexamers consisting of both wild-type and mutant p97 subunits. In such scenario, it will be important to determine if the inhibition of the wild-type subunits in the hexamers by p97 inhibitors can, in fact, compromise the function of the entire hexamer and contribute to sensitivity to p97 inhibitors. CRISPR/Cas9-mediated homology-directed repair can be used to correct the N616fs* mutation in HCT116 resistant cells to answer this important question. Interestingly, in our previous studies, we observed activating mutations (E470D and E470K) in one *p97* allele and inactivating mutations (Q603* and N616fs*) in another *p97* allele in OVSAHO ovarian cancer cells that were selected for resistance to CB-5083 [3, 4]. These results suggest heterozygous mutations in one *p97* allele may not be sufficient to produce resistance to p97 inhibitors unless the other allele is subjected to inactivating mutations or perhaps separate activating mutations.

In summary, we report a previously unidentified missense mutation (L526S) in *p97* that result in resistance towards CB-5083. Moreover, this mutant does not contribute to cross-resistance with NMS-873. More importantly, we observed a pre-existing heterozygous frameshift deletion in *p97* at codon 616 (N616fs*), which help to explain the differences in missense mutation zygosity reported by Anderson *et al.* and Her *et al.*

Although clinical trials of CB-5083 for solid tumors and hematological malignancies were recently prematurely terminated due to off-target effect on PDE6 resulting in ocular dysfunction, additional p97 inhibitors are being explored as clinical leads for cancer treatment. Given that different p97 inhibitors such as CB-5083 and NMS-873 display similar mechanism of drug resistance associated with on-target mutations in p97, it is important to investigate mutations through sequencing of p97 for resistance mechanisms associated with new clinical leads. Such sequencing must be done both at the genomic and complementary DNA level to appropriately delineate the effects of homozygous or heterozygous p97 mutants in drug resistance.

## Materials and Methods

### Cell Lines and cell culture

HCT116 cells (HCT116-par) were purchased from the American Type Culture Condition (ATCC, CCL-247). HCT116-resistant cells (HCT116-res) were generated through ten rounds of intermittent dosing with CB-5083 for 24 hours followed by a recovery phase in the drug-free medium for 5-10 days. CB-5083 treatment started at 2.5 μM and with every round the concentration of CB-5083 was increased by 0.5 μM. Both HCT116-par and HCT116-res were cultured in McCoy’s 5a Medium (ATCC, 30-2007) with 10% fetal bovine serum (Sigma-Aldrich, F8067). Both cells were incubated in a humidified incubator with 5% CO2 at 37^°^C and were checked for mycoplasma contamination.

### Chemicals and cell viability assay

CB-5083 (Selleckchem, S8101) and NMS-873 (Selleckchem, S7285) were purchased from Selleckchem. Both compounds were dissolved in DMSO to make a 50 mM stock solution. Sulforhodamine B (SRB) assay was used to determine the cell viability following the treatment with CB-5083 and NMS-873. SRB assay was performed according to our previously established protocol [7].

### Genomic and complementary DNA sequencing

Genomic and complementary DNA extraction, PCR amplification and sequencing were performed according to our previously established protocols [3]. Information regarding the primers and sequencing results can be obtained in Supplementary 3 and Supplementary 4 respectively.

### Declaration of Interests

Authors have no competing financial interest to disclose.

## Supplementary information

**Supplementary 1:** Mutations in p97 reported in Cancer Cell Line Encyclopedia (CCLE)

**Supplementary 2:** Mutations in p97 reported in The Cancer Genome Atlas (TCGA)

**Supplementary 3:** List of primer sequences.

**Supplementary 4:** Zip Compressed File containing 8 Sanger sequencing results in .ab format.

## References

[1] N.-G. Her, J.I. Toth, C.-T. Ma, Y. Wei, K. Motamedchaboki, E. Sergienko, M.D. Petroski, p97 composition changes caused by allosteric inhibition are suppressed by an on-target mechanism that increases the enzyme’s ATPase activity, Cell chemical biology, 23 (2016) 517–528.

[2] D.J. Anderson, R. Le Moigne, S. Djakovic, B. Kumar, J. Rice, S. Wong, J. Wang, B. Yao, E. Valle, S.K. von Soly, A. Madriaga, F. Soriano, M.-K. Menon, Z.Y. Wu, M. Kampmann, Y. Chen, J.S. Weissman, B.T. Aftab, F.M. Yakes, L. Shawver, H.-J. Zhou, D. Wustrow, M. Rolfe, Targeting the AAA ATPase p97 as an approach to treat cancer through disruption of protein homeostasis, Cancer cell, 28 (2015) 653–665.

[3] P. Bastola, F. Wang, M.A. Schaich, T. Gan, B.D. Freudenthal, T.F. Chou, J. Chien, Specific mutations in the D1-D2 linker region of VCP/p97 enhance ATPase activity and confer resistance to VCP inhibitors, Cell Death Discov, 3 (2017) 17065.

[4] P. Bastola, J. Chien, Co-selected mutations in VCP: a novel mechanism of resistance to VCP inhibitors, Cell Death Dis, 9 (2018) 35.

[5] S.A. Wacker, B.R. Houghtaling, O. Elemento, T.M. Kapoor, Using transcriptome sequencing to identify mechanisms of drug action and resistance, Nature chemical biology, 8 (2012) 235–237.

[6] F. Teraishi, S. Wu, L. Zhang, W. Guo, J.J. Davis, F. Dong, B. Fang, Identification of a novel synthetic thiazolidin compound capable of inducing c-Jun NH2-terminal kinase-dependent apoptosis in human colon cancer cells, Cancer Research, 65 (2005) 6380–6387.

[7] P. Bastola, L. Neums, F.J. Schoenen, J. Chien, VCP inhibitors induce endoplasmic reticulum stress, cause cell cycle arrest, trigger caspase-mediated cell death and synergistically kill ovarian cancer cells in combination with Salubrinal, Molecular oncology, 10 (2016) 1559–1574.

